# Taxonomic Resolution of 16S rRNA, FastANI, Mash, and FastAAI across 30,495 Prokaryotic Type-Strain Genomes

**DOI:** 10.64898/2026.07.15.738563

**Authors:** David Ussery, Momtaz Zamila Bukharid, Ranabir Majumder, Veniamin A Borin, Arghavan Alisoltani

## Abstract

Prokaryotic taxonomy now relies on both marker-gene and genome-wide sequence comparisons, but these methods differ in taxonomic range, scalability, and sensitivity to genome quality. Here, we benchmarked four commonly used approaches, including 16S rRNA identity, FastANI, Mash distance, and FastAAI/Jaccard similarity across a dataset of 30,495 prokaryotic type-strain genomes. Type-strain genomes provide nomenclatural anchors for validly named species, making them a useful framework for evaluating how sequence-based methods correspond to current taxonomic assignments. We evaluated method behavior across taxonomic ranks from species to domain and separated initial method failures from threshold-based failures. When clean full-length 16S rRNA sequences were available, same-species comparisons passed the empirical threshold in >97% of cases. However, a usable full-length 16S rRNA sequence was unavailable for 4,551 of the 16,402 same-species comparisons (28%), limiting marker-gene-based analysis. In addition, 16S rRNA identity ranges overlapped across higher taxonomic ranks, limiting the use of universal rank-specific cutoffs. FastANI provided strong species-level resolution, with same-species comparisons passing the empirical threshold in approximately 88% of cases but was less informative at deeper ranks. Mash enabled rapid genome-scale screening, although its distance values require careful interpretation beyond close relatives. FastAAI provided a genome-wide amino-acid signal, with approximately 92% of same-species comparisons passing the empirical threshold and was especially useful for comparisons beyond the species boundary. Overall, no single method performed optimally across all taxonomic levels. These results support a rank-aware benchmarking framework in which 16S rRNA, FastANI, Mash, and FastAAI are interpreted as complementary tools, with attention to genome quality, missing data, and method-specific failure modes.

## 1. Introduction

Type strains anchor the nomenclature of valid prokaryotic species, with 30,495 bacterial and archaeal type-strain genomes available in NCBI as of May 4, 2026. The tradition of using reference strains stems from early pure-culture studies, such as Koch’s 1876 isolation of *Bacillus anthracis* [1], Escherich’s 1885 description of *Escherichia coli* [2], and Migula’s 1895 structured classification [3]. Many early microbiologists have had genera named after them (**Supplementary Table 1)**. Early bacterial classification based on morphology, physiology, and disease association was formalized in the 1923 first edition of *Bergey’s Manual of Determinative Bacteriology* in 1923 [4]. Bacteria separated from botanical nomenclature under the 1947 Bacteriological Code [5], and type strains remain central to prokaryotic nomenclature today [6], and projects like the Genomic Encyclopedia of Bacteria and Archaea (GEBA) emphasize sequencing these diverse strains [7].

Molecular sequence data transformed prokaryotic systematics. Woese and Fox’s 1977 rRNA comparisons revealed Archaea as a distinct lineage [8], establishing the three-domain framework [9]. Their early, labor-intensive rRNA fingerprinting [8, 10], and subsequent ribosomal studies built the first broad view of microbial evolution [11, 12]. Today, modern sequencing generates complete genomes for thousands of type strains at an unprecedented scale [13, 14]. The broadly conserved 16S rRNA gene is currently the central marker for microbial taxonomy. Historically, a 97% identity threshold was widely adopted for species-level classification [15], though later analyses suggested a bit higher threshold (98.65%) better approximates genome-based boundaries [16]. Today, ≥99% identity is often required for species delineation. Broader rank-associated thresholds have also been proposed [17] and recently re-evaluated using modern taxonomic sampling [18]. Because 16S rRNA identity ranges overlap across ranks, universal cutoffs must be interpreted cautiously. These commonly used thresholds and their overlapping ranges are summarized in **Figure 1**.

**Figure 1.**
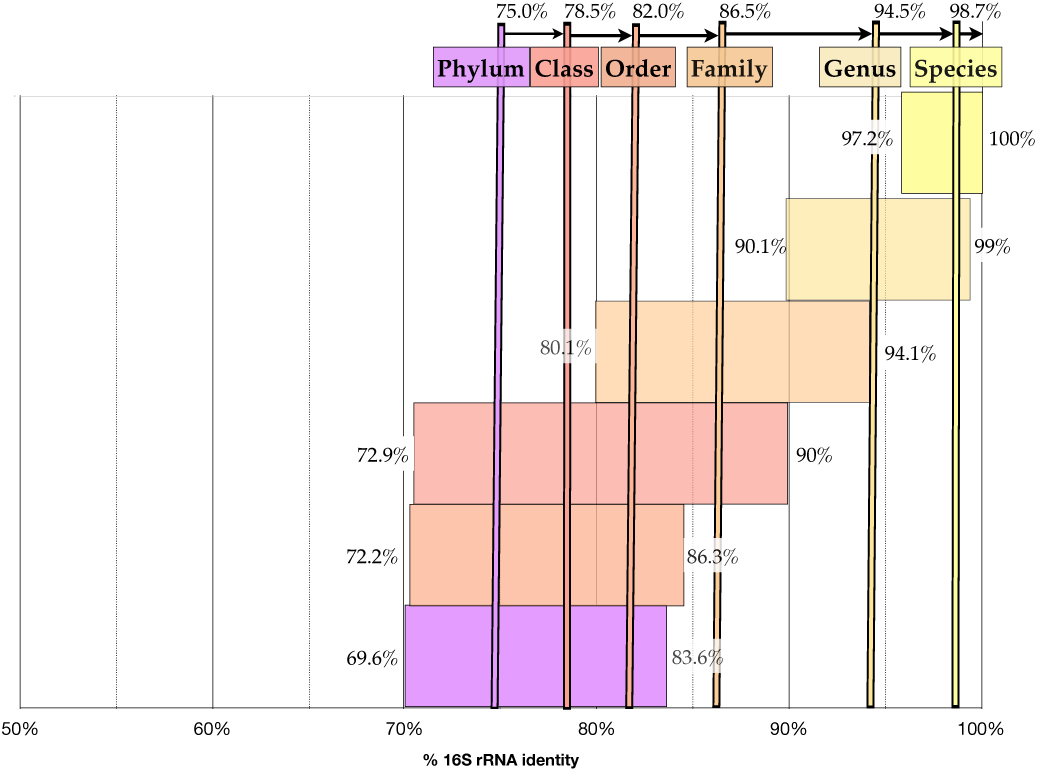
Approximate 16S rRNA identity ranges associated with prokaryotic taxonomic ranks. Vertical markers indicate commonly used rank-associated thresholds reported by Yarza *et al.* [17]. The shaded ranges summarize more recent estimates derived from modern type-strain genome sampling [18]. Because the ranges overlap, the values should be interpreted as empirical guides rather than universal taxonomic boundaries.

Intragenomic 16S rRNA copies are not always identical [19] exhibiting up to 2.4% sequence diversity within some *Vibrio vulnificus* genomes [20] or even more than 5% divergence for environmental adaptation in other organisms [21]. Furthermore, 16S rRNA genes can undergo horizontal transfer [22]. Because this heterogeneity can overestimate genomic diversity [23, 24] and conflict with whole-genome phylogenies [23], modern taxonomy is transitioning toward using full-genome sequences rather than only one or a few marker genes. Based on 16S rRNA, Woese proposed 12 prokaryotic phyla in 1987, and this quickly expanded to 36 [25], then over 50 by 2003 [26], and surpassed 100 following candidate phyla radiation studies (CPR) [27]. The phylum rank remained unofficial until a 2015 proposal [28] was formally adopted by the ICSP and in October 2021, the official list of 42 prokaryotic phyla was validly published [29]. Today, 54 unique phyla are officially recognized by the International Code of Nomenclature of Prokaryotes [30]; the nomenclatural list contains 57 phylum names, including three synonyms [30]. On the NCBI genome web pages, two of these synonyms (*Cyanobacteriophyta* and *Desulfobacterota*) redirect to preferred names, but *Halobacteriota* currently does not point to its synonym, *Methanobacteriota*. In March 2025, NCBI elevated Bacteria and Archaea to Domains, establishing four Bacterial and three Archaeal Kingdoms [31]. Concurrently, the number of proposed phyla has surged. The Genome Taxonomy Database (GTDB) currently lists 186 phyla (R232 [32]), while NCBI data [33] contain 219 genomic prokaryotic phyla (59 Archaeal Phyla [34] and 160 Bacterial Phyla [35]).

Genome sequencing has shifted prokaryotic taxonomy toward genome-wide frameworks like the Genome Taxonomy Database [36]. Despite varying assembly qualities, draft sequencing enables broad gene sampling across genomes [37]. Studies on *E. coli* [38, 39] and other pangenomes [40] reveal massive strain-specific gene pools, underscoring the value of whole-genome comparisons. Average nucleotide identity (ANI) replaced DNA-DNA hybridization [41] and expanded to genome sequences [42, 43], with a 95–96% discontinuity defining species boundaries [43, 44]. Recent guidance based on 330 bacterial and 30 archaeal groups confirms that this ANI gap is common but not universal and should be evaluated within the lineage of interest [45]. Scalable tools such as FastANI [44], EzBioCloud [46], and GTDB [47, 48] have cemented ANI as a species-level standard, though it loses resolution at deeper evolutionary distances. Alignment-free methods like Mash use MinHash sketching to rapidly estimate genome distances within the same species or amongst closely related species [49]. Because these methods differ in their optimal taxonomic ranges and sensitivity to genome completeness, a meaningful benchmark requires a dataset with stable taxonomic anchors, standardized metadata, and sufficient diversity to evaluate performance from species-level comparisons to deeper evolutionary comparisons.

Type-strain genomes are crucial for benchmarking because nomenclature has traditionally been anchored to type material [6, 50]. Type strains are used to evaluate species thresholds [43, 44] and guide GTDB clustering [47]. For comparisons beyond the species level, amino acid methods like FastAAI [51] provide the necessary broader evolutionary signals. Taxonomic instability further necessitates reproducible benchmarking. In 1995, the first bacterial genome was published [52], and the same year, the genome of *Mycoplasma genitalium* was the first bacterial type strain to be sequenced and published. Since then, *M. genitalium*’s classification has shifted across NCBI domains [31] and phyla—from Firmicutes in 1978 [53] to *Tenericutes* in 2008 [54] to *Mycoplasmatota* in 2021 [29]— alongside lower-rank revisions [55]. Only the species name (*M. genitalium*) and ‘Bacteria’ remain, all the intermediate taxonomic ranks have had their names changed. This illustrates why benchmarking studies should use clearly defined genome sets.

Here, we benchmark 16S rRNA, FastANI, Mash, and FastAAI/Jaccard across 30,495 type-strain genomes. Our workflow evaluates these methods from species to domain levels, distinguishing between initial and threshold failures, to offer practical guidance for modern systematics.

## 2. Methods

### Type-strain genomes data retrieval from NCBI

Type-strain genome records were retrieved from the NCBI Datasets web interface on 4 May, 2026 using the “type material only” filter. Bacterial genomes were obtained using the Bacteria query (taxon ID 2) [56], and Archaea (taxon ID 2157) [57]. For each record, the GenBank assembly accession beginning with GCA_ was retained as the primary genome identifier. The bacterial and archaeal accession lists were combined, duplicate GCA accessions were removed, and the resulting frozen dataset contained 30,495 type-strain genomes, comprising 29,383 bacterial and 1,112 archaeal genomes. All accession lists, standardized taxonomy, compact NCBI assembly metadata, analysis and scripts, and supporting documentation for this study are publicly available through the project GitHub repository (https://github.com/ArghavanAlisoltani/30k-type-strain-taxonomy-benchmark).

### Standardizing the taxonomy for the NCBI type-strain genomes

At first, we attempted to use the taxonomy in the headers of the GenBank files. However, we discovered that the taxonomy was changing with time, and some (but not all) of the GenBank files were updated, resulting in an inconsistent taxonomy. To get consistent names, we decided to use the NCBI taxonomy for each of the type strains and made a ‘dictionary’, relating the NCBI Genome Assembly number (GCA_123456789) to the NCBI taxon ID and, from that, extracted the current NCBI taxonomy as of 4 May, 2026. We then checked for missing values in the major taxonomic ranks: domain, kingdom, phylum, class, order, family, genus, and species.

We added missing information as needed (for example, currently NCBI does not list the class for Mollicutes, so we added the same name (“Mollicutes”) as a placeholder to the class for consistency, based on the Genus species name. To do our comparisons, it is necessary to have consistent names across all the taxonomic levels. Each genome assembly accession was used as the main identifier. Blank entries, None, NA, N/A, and nan were treated as missing values. Missing taxonomy was first filled using information already present in the dictionary, where possible. The script created species- and genus-level lookup tables from entries with available taxonomy. If the same species or genus had a consistent assignment elsewhere in the dictionary, that information was used to complete the missing rank.

For entries that could not be completed from the dictionary alone, additional sources were checked, including List of Prokaryotic names with Standing in Nomenclature (LPSN), NCBI, GTDB, Global Biodiversity Information Facility (GBIF), Catalogue of Life, Wikidata, and Wikipedia. Existing taxonomy values were kept unchanged, and only missing fields were filled. After the automated repair step, the completed entries were manually checked for confirmation. We also recorded the source used for each filled value and produced a completed dictionary, a fill report, and a list of entries that still required manual review. The final repaired dictionary was used as the standardized taxonomy reference for downstream genome comparisons.

### Genome Quality Assessment Pipeline

A genome quality control (QC) pipeline was developed to evaluate completeness and sequence integrity across the 30,495 prokaryotic type-strain genomes. The pipeline is based on our previously published quality score approaches [58, 59]. Land *et al*. evaluated genome quality across several public genome repositories, whereas the present study applies an updated version of that framework to a taxonomically anchored collection of 30,495 type-strain genomes and uses the scores to interpret method-specific failures. The four quality scores (sequence, essential genes, rRNA and tRNA) were unchanged with the exception of the rRNA score, which was slightly modified – for this study, we started with 0.1, and added 0.3 for each complete rRNA (5S, 16S, 23S); we gave 0.1 points for a positive score (that is, a gene was found), and another 0.2 points if the full length gene was present (1450-1700 nt for 16S rRNAs). For this project, the presence of full-length 16S rRNA is essential for phylogenetic comparisons, and partial 16S rRNAs were excluded.

### 16S rRNA extraction and pairwise identity analysis

A total of 30,495 publicly available genomes were analyzed using 16S rRNA gene sequences. RNAmmer v1.2 [60] was used with default parameters to identify and extract all predicted 16S rRNA copies from each genome. For the genome-pair benchmarking analysis, one representative full-length sequence per genome was retained, using the highest-scoring prediction when multiple qualifying copies were available. All qualifying copies were retained separately for copy-number and intragenomic heterogeneity analyses. A separate quality-filtered bacterial sequence set was used for the conservation analysis shown in **Figure 6**. This alignment contained 29,438 full-length bacterial 16S rRNA sequences.

The resulting 16S sequences were compiled into a single FASTA file. Pairwise sequence similarity was then computed using VSEARCH (v2.31.0) [61]. An all-versus-all global alignment was performed with the --usearch_global option, using the same dataset as both query and reference database. To ensure exhaustive comparisons, the parameters -- maxaccepts 0 and --maxrejects 0 were applied, and self-hits were included using the --self flag. Sequence identity was calculated using the BLAST-compatible output format (-- blast6out) with identity definition 2 (--iddef 2), which normalizes matches over the full alignment length. No minimum identity threshold was imposed (--id 0.0) to capture the full distribution of similarities.

For selected genome pairs (*e.g.,* predefined taxonomic groupings such as domain or species), percent identity values were extracted by matching query–target genome ID pairs against reference lists. Only the percent identity field (column 3) was retained for downstream analysis. For single pairwise comparisons, only the top hit per query was reported by restricting output to the first record. Summary statistics and distributions of pairwise 16S percent identity were subsequently used as a proxy for genome-level relatedness.

#### Taxonomic Analysis of 16S rRNA Recovery

To assess patterns of 16S rRNA gene recovery across taxonomic levels, a series of post-processing analyses was performed using RNAmmer-generated annotations and genome metadata.

##### Phylum-Level Assessment

Genomes were grouped by phylum based on NCBI taxonomic assignments. For each genome, the corresponding RNAmmer .gff file was queried to identify annotated 16S rRNA features. When present, 16S gene lengths were calculated from start and end coordinates. For each phylum, three quantities were recorded: (i) total number of genomes, (ii) number of genomes containing at least one 16S rRNA gene, and (iii) number of genomes containing at least one high-quality 16S gene with length between 1450 and 1700 bp. The proportion of genomes with high-quality 16S sequences was then calculated per phylum. Results were summarized and sorted by decreasing percentage of high-quality 16S recovery.

##### Genus-Level Missing 16S Analysis

Genomes were also grouped at the genus level to quantify the absence of detectable 16S rRNA genes. An index of RNAmmer .gff files was constructed to map genomes to their annotations efficiently. For each genus with at least five genomes (to reduce sampling noise), the total number of genomes and the number lacking annotated 16S rRNA genes were determined. A missing-16S fraction was calculated as the proportion of genomes within each genus without detectable 16S. Summary statistics included the total number of genera analyzed, the average missing rate, and identification of genera with the highest and lowest missing 16S fractions.

##### Species-Level Representation Analysis

To determine whether missing 16S rRNA genes are associated with undersampled taxa, genomes lacking detectable 16S annotations were further analyzed at the species level. Genome identifiers were standardized and mapped to species names. Species abundance was calculated across the full dataset 30,495 genomes), and each genome was annotated with its corresponding species count. Genomes were classified as single-species genomes if their species was represented by only one genome in the dataset, and multi-species genomes otherwise. This classification enabled comparison of missing 16S incidence between rare (single-representation) and well-sampled species.

### Intragenomic 16S rRNA heterogeneity analysis

For each bacterial species, two representative genomes were selected from NCBI using genome accession identifiers (GCA). Each pair consisted of one designated type-strain genome and one additional representative genome from the same species, ensuring consistent taxonomic comparison across datasets.

#### Identification of 16S rRNA sequences

Putative 16S rRNA gene copies were identified in each genome using RNAmmer (v1.2) [60]. The resulting predictions included genomic coordinates and extracted 16S rRNA sequences per genome.

#### Intragenomic 16S similarity analysis

To quantify intragenomic heterogeneity of 16S rRNA gene copies, pairwise sequence comparisons were performed within each genome using VSEARCH (v2.31.0) [61] with global alignment (--usearch_global). Each 16S sequence was compared against all other 16S copies within the same genome. Self-comparisons were excluded from downstream statistics.

#### Summary statistics

For each genome, the following metrics were computed from pairwise identity distributions:

n: number of pairwise VSEARCH comparisons
min identity: lowest pairwise 16S identity (used as the “worst-case” divergence metric)
avg identity: mean pairwise identity across all comparisons
max identity: highest observed pairwise identity

#### Visualization

For each species, the minimum pairwise identity (worst-case 16S divergence per genome) was visualized as a histogram/bar representation. The type strain genome was explicitly included for comparison against non-type representatives.

### Genome-wide pairwise comparison methods

Genome pairs were assigned to mutually exclusive taxonomic-rank categories using the standardized taxonomy. The species category contained genomes assigned to the same species. The genus category contained genomes assigned to the same genus but different species, and each successively higher-rank category contained genomes sharing the indicated rank while differing at all lower evaluated ranks. The same set of genome-pair identifiers was used to integrate the 16S rRNA, FastANI, Mash, and FastAAI/Jaccard results.

FastANI v1.33 [44] was used to calculate average nucleotide identity for the predefined genome pairs. Genome assemblies were supplied as query and reference FASTA files, and comparisons were performed using default parameters. The reported ANI percentage was retained for downstream analysis. A pair was classified as an initial failure when FastANI did not return a valid ANI estimate. Among valid same-species comparisons, values below 96.94, that is mean minus one standard deviation (SD), were classified as threshold failures.

Mash v2.3 [49] was used to estimate genome distances using MinHash sketches. Genome assemblies were sketched using a k-mer size of 21 and a sketch size of 1000, and pairwise distances were calculated using `mash dist`. Mash distance ranges from zero for highly similar sketches to one as detectable similarity decreases. A pair was classified as an initial failure when no valid Mash distance was returned. Among valid same-species comparisons, distances greater than 0.04, that is mean *plus* one SD, were classified as threshold failures.

FastAAI v0.1.17 [51] was used to calculate amino-acid-based genome similarity using tetramer profiles derived from universal proteins. Databases and pairwise comparisons were generated using default options. The FastAAI/Jaccard similarity value was retained for the rank-distribution and same-species analyses, with higher values indicating greater similarity. A pair was classified as an initial failure when FastAAI did not return a valid similarity value. Among valid same-species comparisons, similarities below 0.91 (mean minus one SD) were classified as threshold failures. For Neighbor-joining tree construction, FastAAI/Jaccard similarity was converted to distance as 1 − Jaccard similarity.

Initial failures and threshold failures were treated as separate outcomes. Initial-failure percentages were calculated using all 16,402 same-species comparisons, whereas threshold-failure percentages were calculated using only comparisons for which the corresponding method produced a valid score.

### Phylum-level 16S rRNA gene tree and FastAAI trees

For the phylum-level tree analysis, we sampled one representative genome per phylum along with the genome for *Microcaldota archaeon* (accession GCA_021654395) as an outgroup to ensure comprehensive taxonomic coverage. Consequently, this additional genome is unique to the phylogenetic analysis and was not included in the broader 30,495-genome benchmark summaries, assembly-status distributions, and method-performance evaluations. Representative full-length 16S rRNA sequences were aligned with MAFFT (v7.475) [62], and a maximum-likelihood tree was inferred with IQ-TREE (v 2.0.7) [63] using its Model Finder function [64] and ultrafast bootstrap of 1000 replicates. The resulting tree was visualized using custom Python code, with concentric annotation rings indicating Domain, Kingdom, and Phylum assignments from the standardized NCBI taxonomy table.

Within the reduced phylum-representative dataset used to construct Figure 9, ten genomes contained more than one retained full-length 16S rRNA copy. Pairwise nucleotide distances for these ten genomes were calculated as 1 − pairwise identity from the MAFFT alignment and are summarized in Supplementary Table 2. This targeted subset was derived exclusively from the **Figure 9** tree dataset and was independent of the full 30,495-genome copy-number analysis shown in **Figure 4**. To compare the phylum-level signal recovered by the reduced 16S rRNA phylogeny and the FastAAI Neighbor-joining tree, one of the longest qualifying 16S rRNA copies was retained per genome after confirming that within-genome copies were more similar to one another than to external representatives. The FastAAI Neighbor-Joining and 16S rRNA trees were pruned to the same genome set and rooted using the archaeal genomes as a shared outgroup to ensure that both trees received the same directional interpretation. Tree agreement was evaluated using topology-based metrics, including Robinson–Foulds (RF) distance, quartet distance, and rooted cluster distance; branch-length-aware agreement was assessed using Mantel correlations of pairwise patristic-distance matrices. A tanglegram was then generated after leaf-order optimization to visualize taxon-level concordance between the two rooted trees. Tanglegram construction, tree-order optimization, tree-distance calculations, Mantel tests, and figure generation were performed using custom Python v3.13.13 scripts. Numerical and tabular operations used NumPy v2.4.6 and pandas v3.0.3, respectively, and figures were generated using Matplotlib v3.10.9. Tree parsing, rooting, pruning, branch-length handling, Robinson–Foulds distance, rooted cluster distance, weighted branch-score distance, quartet distance, patristic distance calculations, and Mantel permutations were implemented directly within the custom Python workflow available in the aforementioned GitHub repository.

## 3. Results

### Overview of the 30,495 NCBI type-strain genomes

The explosion in bacterial genome sequences in GenBank is shown in **Figure 2A**, along with the ratio of complete to draft genomes. The insert shows that by 2010, about half of the genomes were draft. Currently, around 97% of the bacterial genomes in GenBank are draft, and only 3% are complete. For this project, this means that many bacterial genomes lack complete, full-length 16S rRNA, making the use of this gene for taxonomic purposes difficult. Fortunately, advances in long-read, single-molecule third-generation sequencing have enabled more complete genomes to be deposited in GenBank. For example, out of frustration with the lack of a finished type strain for *E. coli*, several years ago, we sequenced the complete *E. coli* type strain in a few days, for less cost than the open access journal publication fees [65].

**Figure 2.**
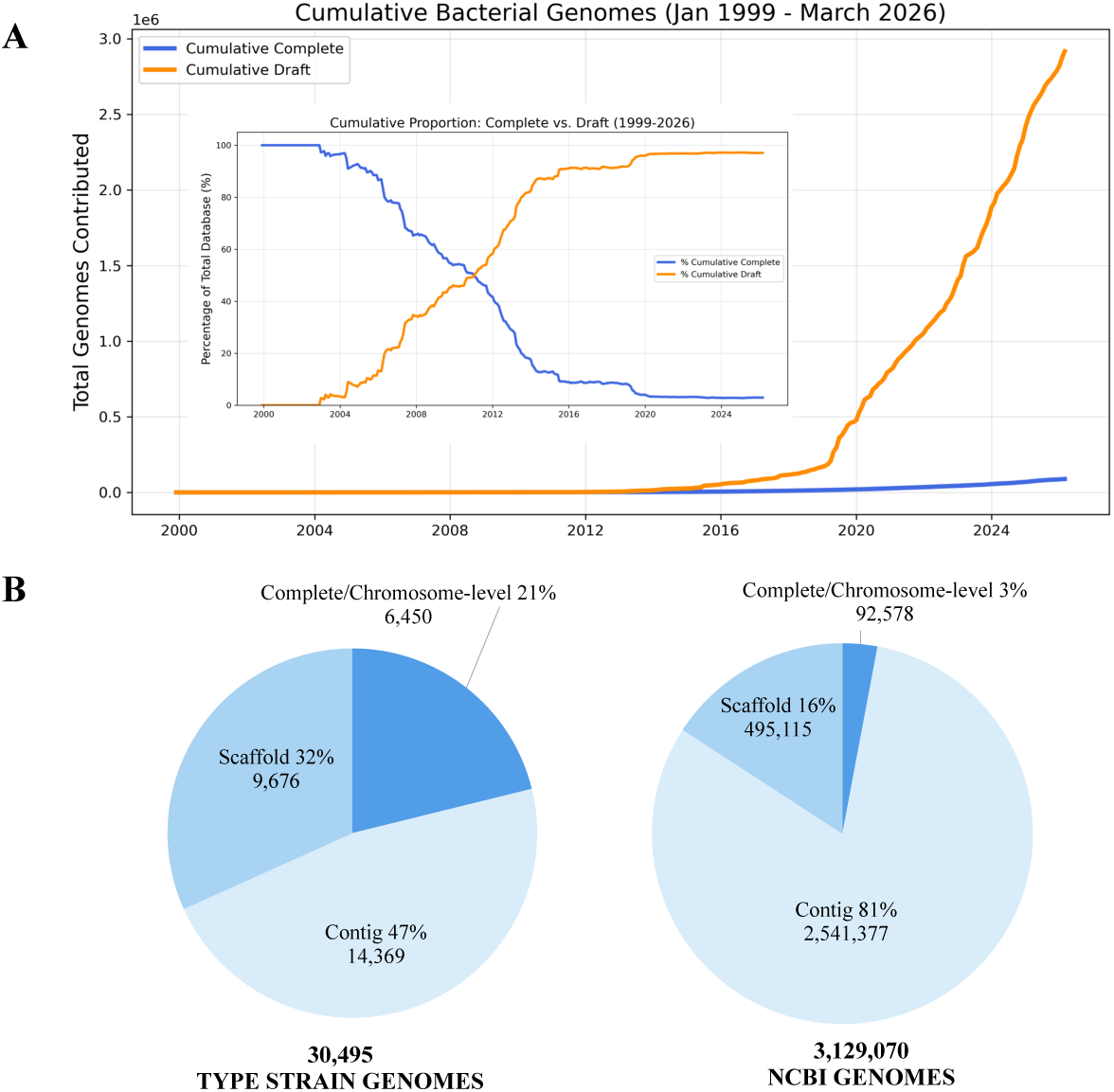
(A) Bacterial genome sequencing growth (main plot) and quality (insert) over the past 30 years. (B) Assembly composition of bacterial genomes from NCBI. The pie chart shows the proportion of genomes classified as complete/chromosome-level, scaffolds, or contigs, as defined by NCBI. Complete represents fully assembled genomes; scaffolds are draft collections of ordered contigs; and contigs represent unjointed sequence fragments (lowest quality). Assembly completeness of bacterial type strains (left) compared to the ∼3 million total bacterial genomes in GenBank as of March 2026 (right).

The type-strain genome set was of higher assembly quality than the broader NCBI genome collection, but draft assemblies still dominated: 6,450 genomes (21.2%) were complete or chromosome-level, 9,676 (31.7%) were scaffold-level, and 14,369 (47.1%) were contig-level assemblies. **Figure 2B** shows the relative proportions of quality in type-strain genomes compared to the more than 3 million bacterial genomes currently in GenBank.

### Taxonomic Distribution of NCBI type-strain genomes

We retrieved all the prokaryotic type strain genomes from NCBI, which at the time (May, 2026) was 30,495 genomes. As shown in **Figure 3**, the dataset contained two domains, Archaea and Bacteria, distributed across nine NCBI kingdoms. According to NCBI taxonomy, the collection of genomes from type material represented a total of 2 Domains, 9 Kingdoms, as shown in **Figure 3**. The downloaded genomes were from 88 Phyla, 197 Classes, 431 Orders, 1003 Families, 4470 Genera, and 21,971 unique species. Notice that, even though these 30,495 type strains are supposed to represent diverse phylogeny, in practice only a few taxonomic groups dominate, with a typical power-law distribution, as shown in **Supplementary Figure 1**. The type strains that are easiest to grow and work with in the lab are often over-represented, and environmental samples that require more special growth requirements can be under-represented.

**Figure 3.**
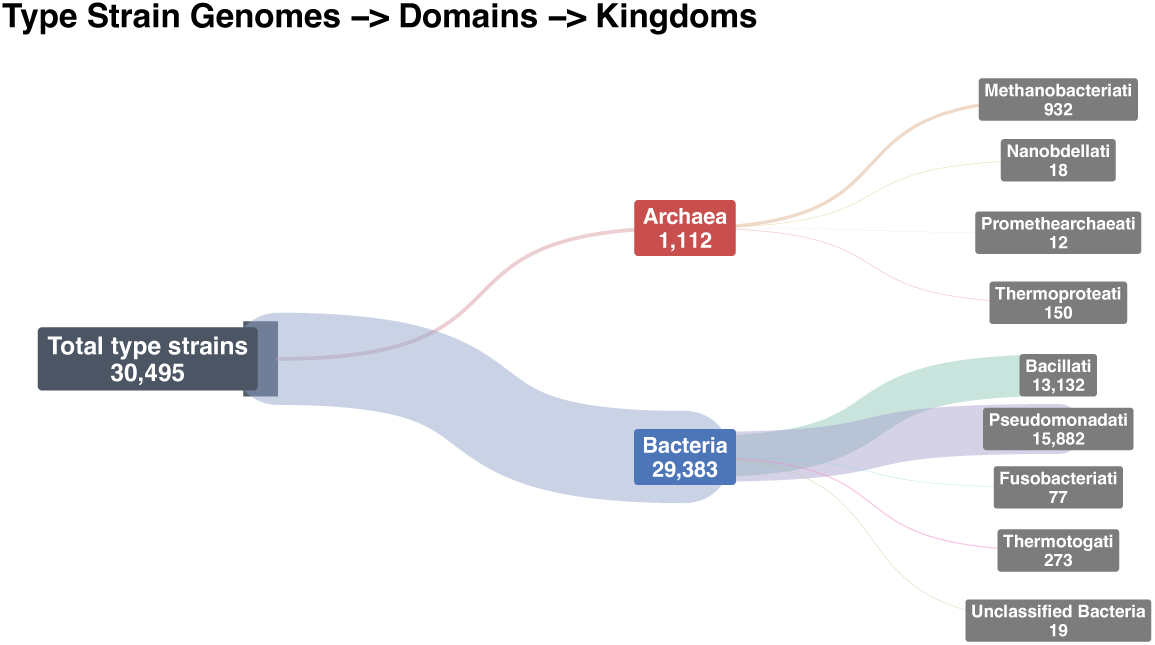
Distribution of type strains across taxonomic groups. Note that in Archaea one Kingdom dominates (*Methanobacteriati*), and that in Bacteria only two Kingdoms (*Bacillati* and *Pseudomonadati*) account for most [>95%] of the NCBI type strain genomes.

The 30,495 type-strain genomes represented 21,971 species. Most species were represented by very few genomes: 21,346 species (97.2%) had three or fewer genomes, and 16,584 species (75.5%) were represented by a single genome (**Table 1**). Only 27 species were represented by ten or more genomes. The most highly represented species were *Salmonella enterica* with 38 genomes (for historical reasons, there are multiple *S. enterica* type strains), *Bacillus subtilis* with 25, and *Lactobacillus delbrueckii* and *Pseudomonas chlororaphis* with 22 each. This uneven species-level sampling influences the number of available same-species comparisons and emphasizes the importance of distinguishing genome counts from pairwise-comparison counts.

**Table 1.** Number of genomes for type strain species.

| # genomes | # species | # same species comparisons | Total genomes | # genomes | # species | # same species comparisons | Total genomes |
| --- | --- | --- | --- | --- | --- | --- | --- |
| 1 | 16,584 | 0 | 16,584 | 11 | 5 | 275 | 55 |
| 2 | 3,706 | 3,706 | 7,412 | 12 | 2 | 132 | 24 |
| 3 | 1,056 | 3,168 | 3,168 | 14 | 1 | 91 | 14 |
| 4 | 327 | 1,962 | 1,308 | 15 | 2 | 210 | 30 |
| 5 | 141 | 1,410 | 705 | 17 | 2 | 272 | 34 |
| 6 | 53 | 795 | 318 | 18 | 2 | 306 | 36 |
| 7 | 32 | 672 | 224 | 20 | 1 | 190 | 20 |
| 8 | 29 | 812 | 232 | 22 | 2 | 462 | 44 |
| 9 | 16 | 576 | 144 | 25 | 1 | 300 | 25 |
| 10 | 8 | 360 | 80 | 38 | 1 | 703 | 38 |
|  |  |  |  | Sum | 21,971 | 16,402 | 30,495 |

### 16S rRNAs in the type strain genomes

**Figure 4** plots the copy number of 16S rRNAs per genome for the 30,495 type strain genomes. The average type strain genome for both Archaea and Bacteria has only one copy of the 16S rRNA gene. The average for more than 3 million NCBI complete bacterial genomes contains about 7 copies of the rRNA gene (data not shown; see **Supplementary Figure 2A**), which is likely reflective of the strong bias of “*E. coli* and friends” bacterial species that dominate GenBank). The Ribogrove [66] website contains a count of the average 16S rRNAs, somewhat normalized, with an average 16S rRNA genome copy number per species, and is shown in **Supplementary Figure 2B**. It is also worth noting in **Figure 4** that many of the type strain genomes (5,925 or roughly 20%) have zero copies - that is, they are missing full-length 16S rRNAs.

**Figure 4.**
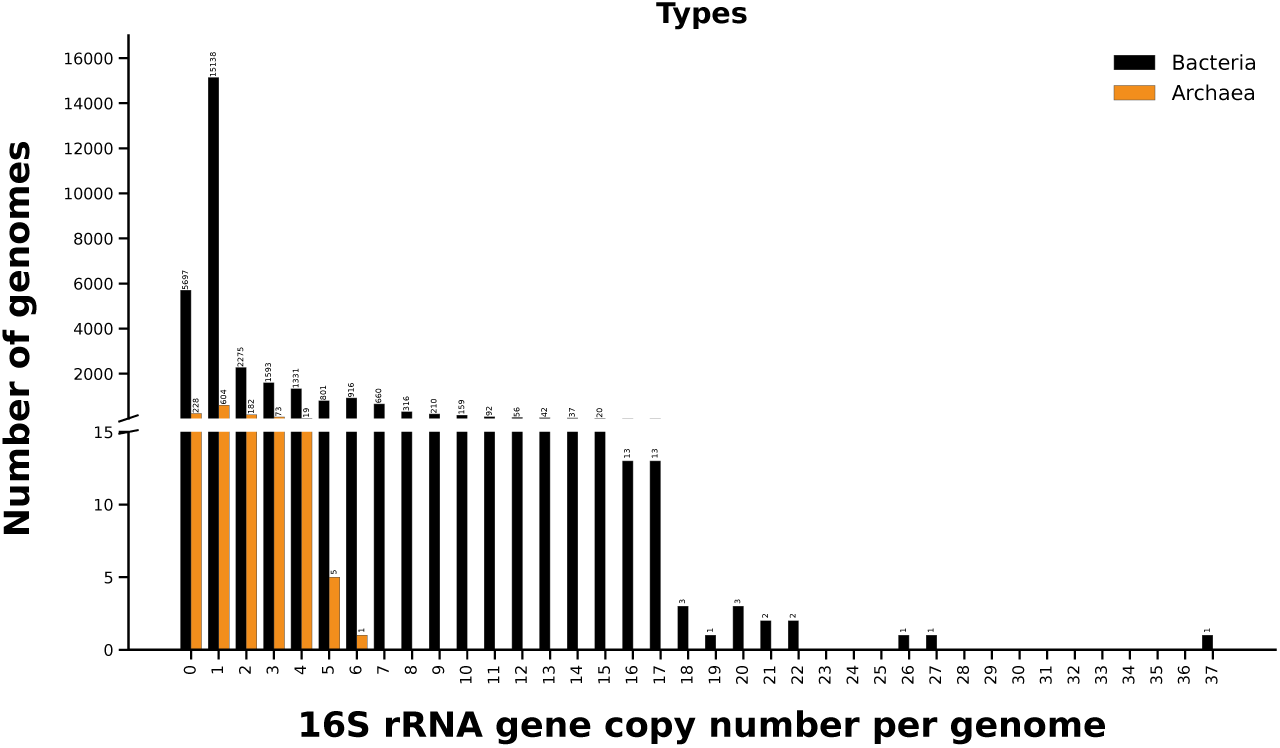
Frequency bar plot showing the number of full-length 16S rRNA gene copies detected per genome (Archaea = Orange; Bacteria = Black). Note that 5,697 bacterial genomes are missing at least one full-length copy of the 16S rRNA gene.

High 16S rRNA copy numbers are uncommon. Sixty genomes contained at least 15 full-length copies, 40 contained more than 15 copies, and ten contained at least 20 copies. The maximum observed copy number was 37 in the *Tumebacillus avium* genome. For the type strain genomes, it is reported that a few genomes [37] have 20 or more copies of the 16S rRNA genes, and more than 100 type-strain genomes contained more than 15 full-length copies of the 16S rRNA gene; 15 copies have been previously reported as the upper limit [67, 68]. However, it appears that most of these genomes with 15 or more copies are complete and well assembled, and it is likely that these high 16S copy numbers are real, although the biological reason is not clear. The genome with the most full-length copies of the 16S rRNA genes was the *Tumebacillus avium* genome, with 37 nearly identical copies of 16S rRNA genes. However, this species divides slowly, taking 2 days, although, as mentioned in the introduction, there is a strong correlation between rRNA copy number and rapid genome division. Previous work has shown that 16S rRNA variation is common in genomes with about a dozen or fewer copies of 16S rRNA genes, whereas genomes with 14 or more copies tend to show little variation within the 16S rRNA genes [23].

### Missing 16S rRNAs in the type strain genomes

**Figure 5** integrates the taxonomic hierarchy of the 30,495 genomes with phylum classification, full-length 16S rRNA availability, and method-specific threshold failures. The dataset contained 26 archaeal and 62 bacterial phylum labels. Seventeen archaeal and 14 bacterial phyla were classified as candidate, unclassified, or other unofficial groups. Thirteen of the 26 archaeal phyla contained no genome with a recoverable full-length 16S rRNA sequence; however, many of these phyla were represented by only one or a few genomes, and percentages of 0% or 100% should therefore be interpreted cautiously. **Supplementary Figure 1A** and **Supplementary Figure 1B** provide the corresponding domain-to-kingdom-to-phylum flows, lower-rank diversity, and genome counts for Archaea and Bacteria.

**Figure 5.**
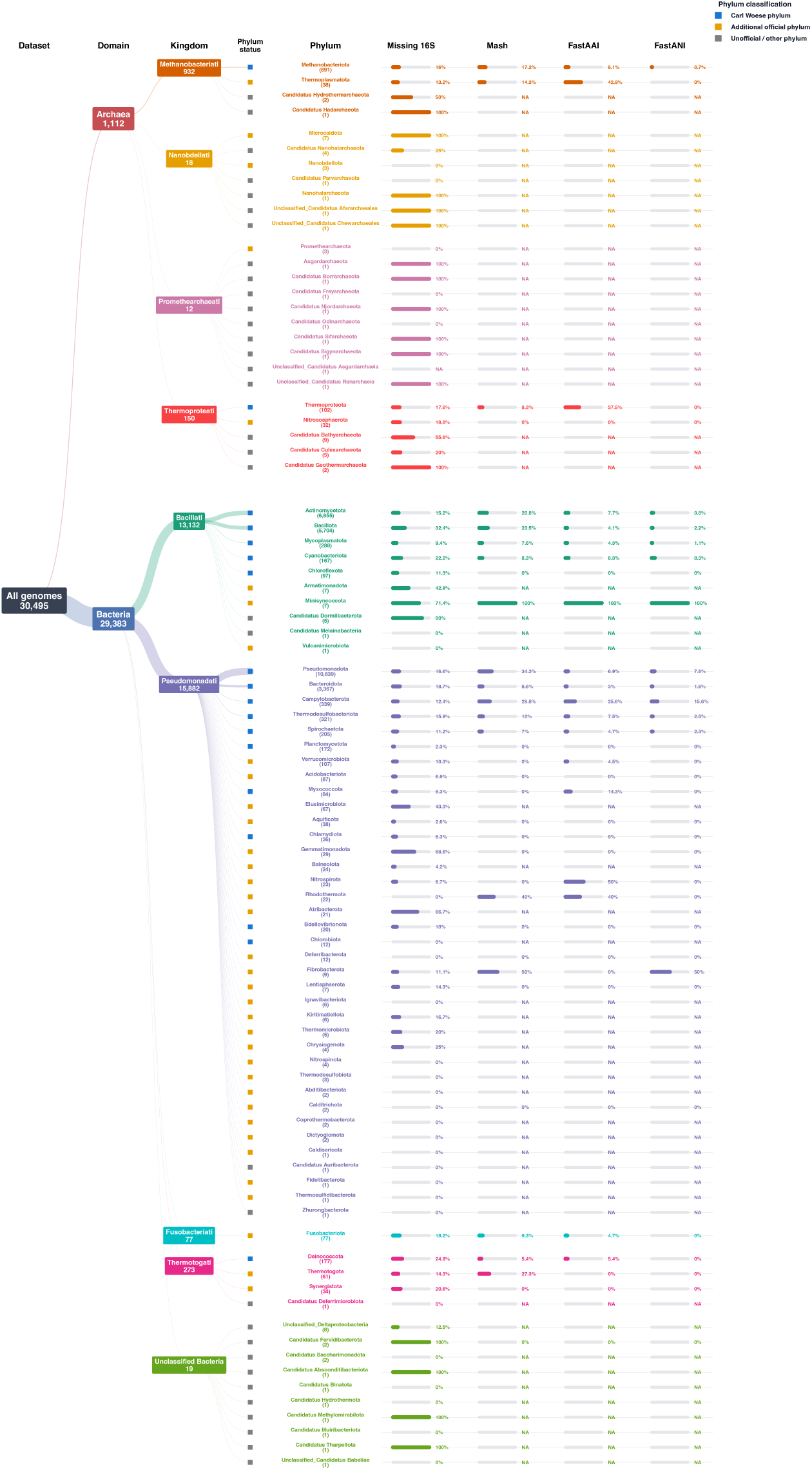
Phylum-level taxonomic structure, full-length 16S rRNA availability, and method-specific threshold failures. Sankey flows trace the 30,495 genomes through domain, kingdom, and phylum. Colored markers classify phyla as corresponding to an original Woese group, an additional officially recognized phylum, or an unofficial/candidate/unclassified group. The Missing 16S column shows the percentage of genomes in each phylum lacking a recoverable full-length 16S rRNA sequence. The Mash, FastAAI/Jaccard, and FastANI columns show the percentage of valid same-species comparisons within each phylum that failed the corresponding empirical threshold. N/A indicates that no eligible same-species comparison was available.

The standardized NCBI taxonomy contained 88 phylum labels, whereas 54 unique phyla were classified as officially recognized in the present analysis. These 54 phyla contained 30,431 of the 30,495 genomes (99.8%), leaving only 64 genomes (0.2%) assigned to candidate, unclassified, or other phylum labels. **Supplementary Figure 2** divides the recognized phyla into 18 phyla corresponding to the original Woese groupings, which together contained 29,676 genomes (97.3% of the dataset), and 36 additional recognized phyla containing 755 genomes (2.5%). Thus, restricting broad comparisons to recognized phyla substantially simplifies the taxonomy while retaining nearly the complete benchmark dataset.

### 16S rRNA alignment across 29,438 bacterial type-strain genomes

To visualize conservation and variability across the marker used for 16S-based comparisons, we examined a quality-filtered alignment of 29,438 full-length bacterial 16S rRNA sequences recovered from type-strain genomes. The alignment includes more than one sequence from genomes with more than one retained copy. Shannon entropy was converted to information content in bits (2-H) and summarized using a 12-nucleotide sliding window. **Figure 6** shows the characteristic mosaic structure of the 16S rRNA gene: highly conserved positions occur throughout the alignment, whereas reductions in information content correspond broadly to the recognized V1–V9 variable regions. Conserved regions facilitate alignment across distantly related bacteria, whereas variable regions provide much of the signal used to distinguish more closely related taxa. However, the uneven distribution of variation and the occurrence of nearly identical 16S rRNA sequences in distinct species limit the use of universal identity thresholds. This analysis extends the original RNAmmer information-content analysis, which used 743 curated bacterial sequences, to a much larger collection of type strains [60]. Despite the substantial increase in taxonomic sampling, the broad pattern of conserved and hypervariable regions remained similar, supporting the robustness of the earlier observations while providing a finer estimate of site-specific 16S rRNA diversity.

**Figure 6.**
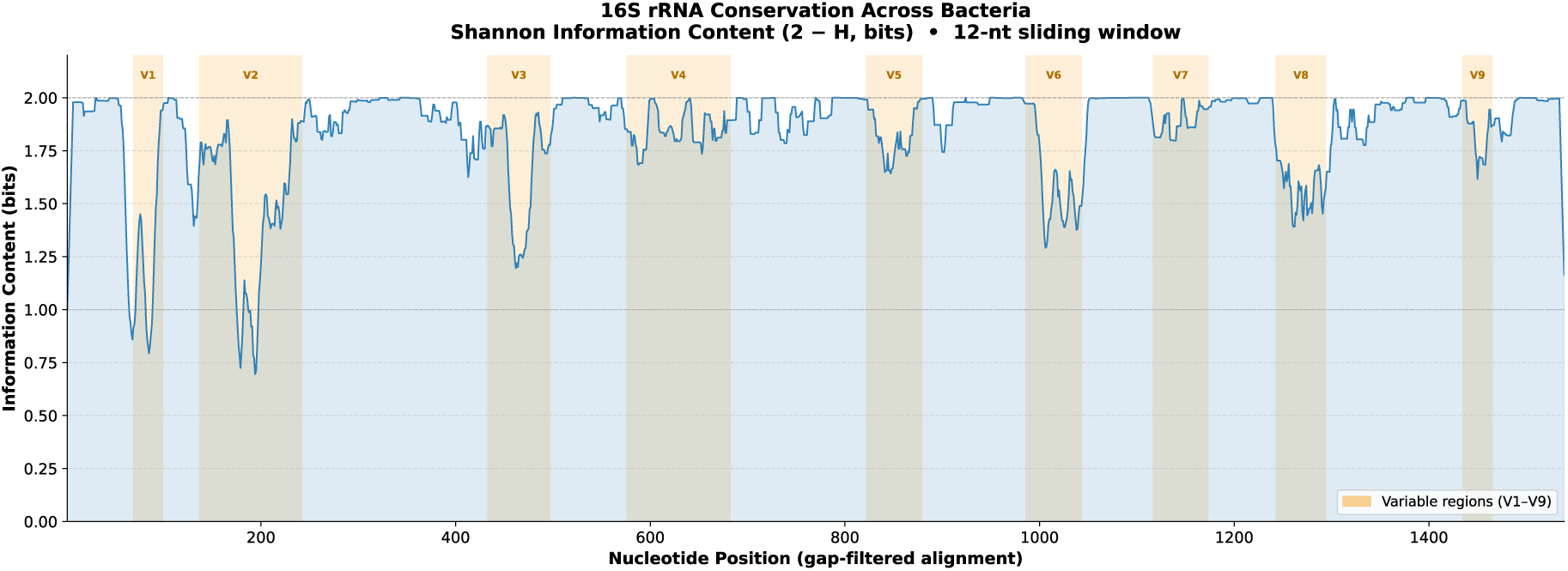
Conservation and variability across the 16S rRNA gene alignment. The alignment highlights conserved and variable regions across full-length 16S rRNA sequences from type-strain genomes.

### Benchmarking 16S rRNA and genome-wide methods across taxonomic ranks

**Figure 7** shows the relative performance of four different methods for identifying taxonomic groups based on their genome sequences. Notice that in the first three methods (**Panels A-C** in **Figure 7**), the ranges only use values in the upper one-third of the scale, and in the fourth method (Jaccard similarity from FastAAI), the full range from 0 to 1 is used. The 16S rRNA comparisons (**Panel A** in **Figure 7**) reveal that most sequence pairs fall within the 70–100% identity range. For this method, a result is always obtained when a 16S gene is present, in contrast to Mash, FastANI, and FastAAI, which may fail to produce a similarity score depending on parameter settings. This illustrates the broad but limited resolution of 16S across taxonomic levels.

**Figure 7.**
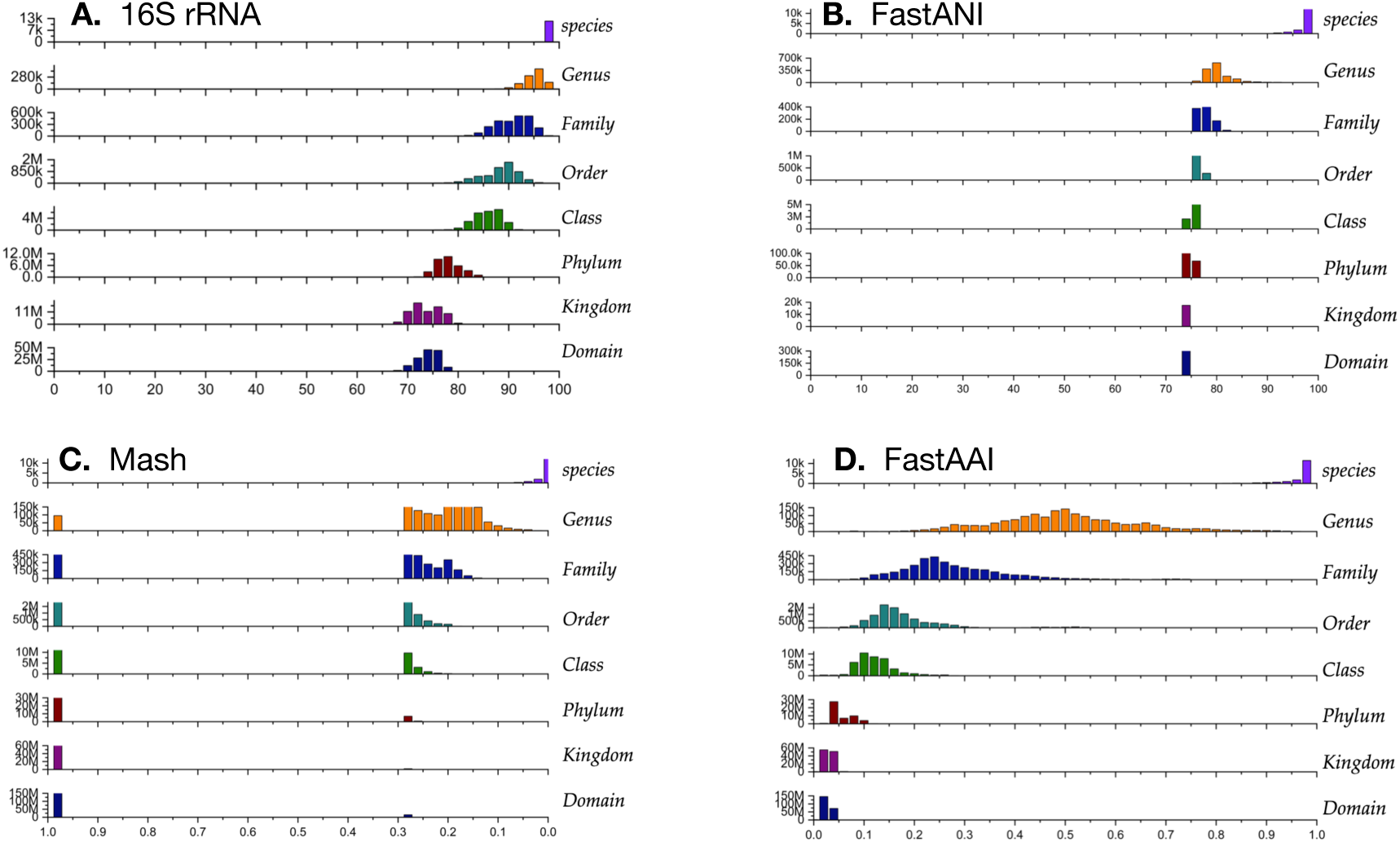
Comparison of four sequence-based methods across taxonomic levels. Pairwise comparisons were grouped by standardized NCBI taxonomic rank. The species row includes genome pairs assigned to the same species among species represented by two or more genomes. The genus row includes pairs from the same genus but different species; higher-rank rows similarly compare genomes sharing the indicated rank while excluding pairs already grouped at lower ranks. **Panel A** shows 16S rRNA percent identity for genome pairs with usable full-length 16S rRNA sequences. **Panel B** shows FastANI values for genome-pair comparisons. **Panel C** shows Mash distances; because Mash is a distance metric, lower values indicate greater similarity, with zero representing identical or nearly identical sketches and larger values indicating decreasing detectable similarity under the chosen Mash parameters. **Panel D** shows FastAAI/Jaccard similarity across taxonomic levels. For 16S rRNA, FastANI, and FastAAI/Jaccard similarity, higher values indicate greater similarity; for Mash distance, lower values indicate greater similarity. Note that each method has a different numerical range and should be interpreted according to its own scale.

FastANI results (**Figure 7, Panel B**) demonstrate very high resolution at the species level, with most comparisons at or near 100% similarity. Genus-level comparisons provide the broadest distribution, indicating maximal resolving power at this taxonomic scale. At higher levels (family, order, class, phylum, domain), the distributions gradually sharpen near ∼70% similarity, while the proportion of comparisons with undetectable similarity (0%) increases, reflecting more distant relationships.

Mash distances (**Figure 7, Panel C**) showed the expected increase with taxonomic separation. Values near zero indicated highly similar genomes, whereas a value of one represented no detectable similarity under the selected sketching parameters. Same-species comparisons were concentrated near zero, while genus- and family-level comparisons covered broader intermediate ranges. At higher ranks, an increasing fraction of comparisons accumulated at one, demonstrating that Mash is most informative for rapid screening of close and moderately related genomes rather than for reconstructing deep taxonomic relationships.

Finally, we used FastAAI, a relatively new method published in 2025 [52]. FastAAI was developed to address two major challenges: the poor scalability of many exhaustive pairwise genome-comparison approaches and the declining resolution of nucleotide-based methods across deeper evolutionary distances. Recent guidance also emphasizes that 16S rRNA identity does not consistently resolve species, because genomes sharing more than approximately 98.5% 16S rRNA identity may belong either to the same or to different species [45]. This agrees with the substantial overlap observed among the species- and genus-level 16S distributions in our analysis. FastAAI complements these approaches by providing a scalable amino-acid-based similarity measure that retains a broader signal across genus, family, order, and deeper comparisons. The FastAAI/Jaccard results are shown in **Figure 7, Panel D**.

**Table 2** summarizes two distinct failure categories among the 16,402 same-species genome comparisons. An initial failure occurred when a method did not produce a usable score, such as when at least one genome lacked a recoverable full-length 16S rRNA sequence. A threshold failure occurred when a valid score was produced but did not meet the empirical same-species threshold defined for that method. Initial-failure percentages were calculated using all 16,402 comparisons as the denominator, whereas threshold-failure percentages were calculated only among comparisons that produced valid scores. These categories were kept separate because they represent different technical and biological limitations.

**Table 2.**
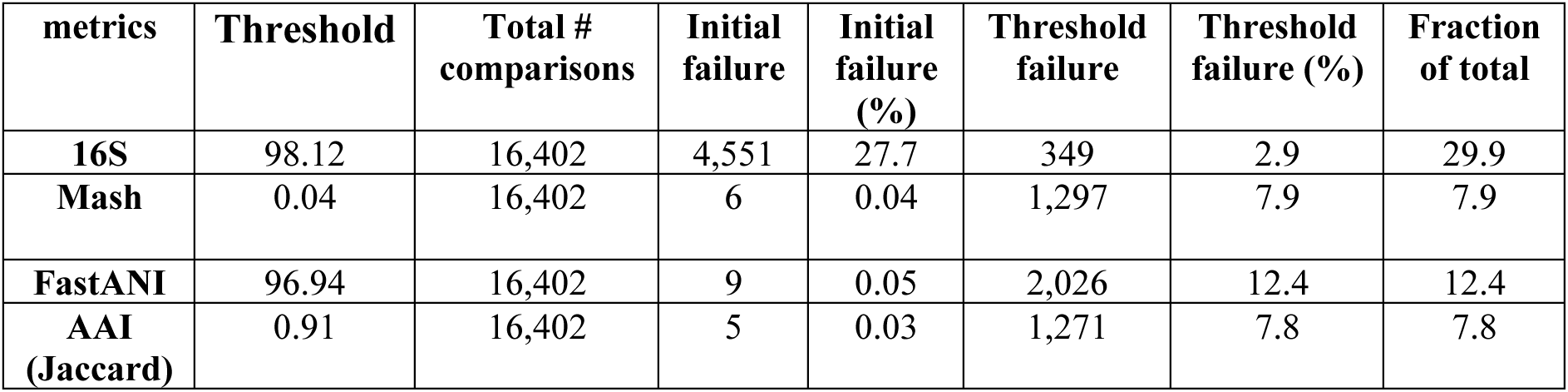
Comparison of initial and threshold failure rates across the four methods. Method-specific thresholds were defined using the mean ± standard deviation, with the direction of each cutoff determined by the scale and interpretation of the corresponding metric. Threshold-failure rates were calculated among valid comparisons. Additional details are provided in the Methods section, “Genome-wide pairwise comparison methods.

The largest initial-failure category occurred for 16S rRNA: 4,551 comparisons (27.7%) could not be evaluated because one or both genomes lacked a usable full-length sequence. Among the remaining 11,851 valid 16S comparisons, only 349 (2.9%) failed the empirical threshold, corresponding to a 97.1% passing rate. Initial failures were rare for the genome-wide methods, occurring in six Mash, nine FastANI, and five FastAAI/Jaccard comparisons. FastANI had the highest threshold-failure rate among valid genome-wide comparisons at 12.4%, whereas Mash and FastAAI/Jaccard had similar rates of 7.9% and 7.8%, respectively. FastAAI/Jaccard therefore provided the lowest combined initial and threshold failure rate among the genome-wide approaches while retaining useful resolution across broader taxonomic distances.

Quality-score distributions for the complete set of 30,495 type-strain genomes are shown in **Supplementary Figure 3**. Scores were concentrated near the upper end of the scale, particularly for the tRNA and Pfam essential-gene components, whereas sequence and rRNA scores showed broader lower tails. This pattern is consistent with the earlier analysis of approximately 32,000 public genomes by Land *et al.* [59], which found generally high combined quality scores but identified sequence and rRNA completeness as important sources of variation. The quality scores were used descriptively to characterize the benchmark dataset and were not used as general exclusion criteria.

Whereas **Table 2** summarizes the total initial and threshold failures for each method independently, **Figure 8** uses exact pair identifiers to show how the threshold-failure sets overlap among methods. Among the 16,402 same-species genome comparisons, 2,815 unique pairs failed the threshold for at least one method. The largest mutually exclusive groups comprised pairs failing both Mash and FastANI only (683 pairs), FastANI only (606 pairs), FastAAI/Jaccard only (561 pairs), and Mash, FastANI, and FastAAI/Jaccard together (517 pairs). Only 61 pairs failed the thresholds across all four methods, representing 0.37% of the complete same-species comparison set. These results indicate that most threshold failures were method-specific or shared among a subset of methods rather than consistently observed across all four approaches.

**Figure 8.**
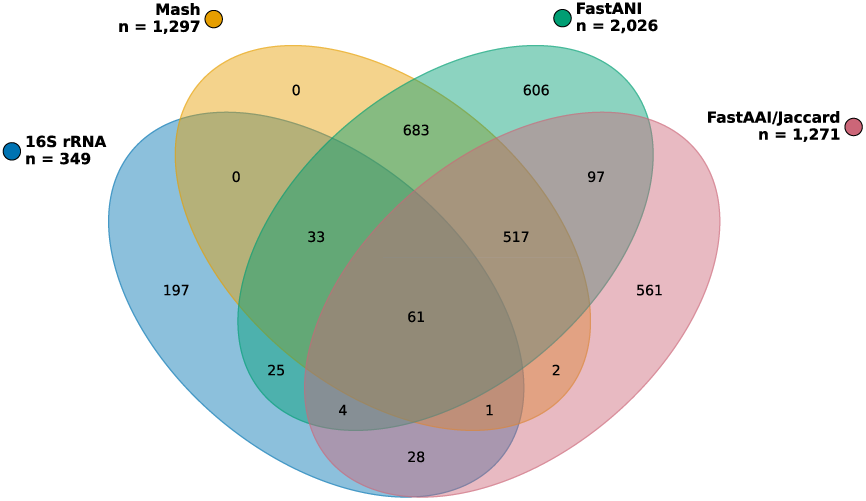
Exact overlap of species-level threshold failures among the four sequence-based methods. Each ellipse represents the same-species genome pairs that produced a valid comparison but failed the corresponding method-specific threshold. The totals shown beside the method names indicate the complete threshold-failure set for each method: 349 for 16S rRNA, 1,297 for Mash, 2,026 for FastANI, and 1,271 for FastAAI/Jaccard. Numbers within the diagram represent mutually exclusive genome-pair intersections calculated from 16,402 exact pair records. Sixty-one pairs failed the thresholds for all four methods. Initial failures were excluded, and ellipse areas are schematic rather than proportional to the counts.

Together, these results illustrate the complementary strengths and limitations of the four methods. When a full-length sequence was recoverable, 16S rRNA produced the lowest threshold-failure rate but showed substantial overlap among closely related taxonomic ranks. FastANI provided strong species-level resolution but had the highest threshold-failure rate among the valid genome-wide comparisons. Mash offered rapid screening of close relationships but progressively lost detectable similarity across deeper ranks. FastAAI/Jaccard combined a low initial-failure rate with broad amino-acid-based resolution across both shallow and deeper taxonomic comparisons.

### Comparison of 16S rRNA gene tree and FastAAI trees of phylum representatives

We further investigated the resolution of the two most informative deep-rank methods—16S rRNA sequence identity/phylogeny and FastAAI Jaccard-distance Neighbor-Joining tree for evaluating phylum-level structure and concordance between marker-gene and genome-wide signals. To evaluate the placement of selected genomes with intragenomic 16S rRNA heterogeneity in a broader taxonomic context, we constructed a 16S rRNA marker-gene tree using representative full-length 16S rRNA sequences from a representative type strain from each of the 56 officially recognized prokaryotic phyla (**Figure 9**). The tree recovered a broad separation between Archaeal and Bacterial representatives, and the taxonomic annotation rings showed that most sequences were placed consistently with their assigned NCBI domain, kingdom, and phylum labels.

**Figure 9.**
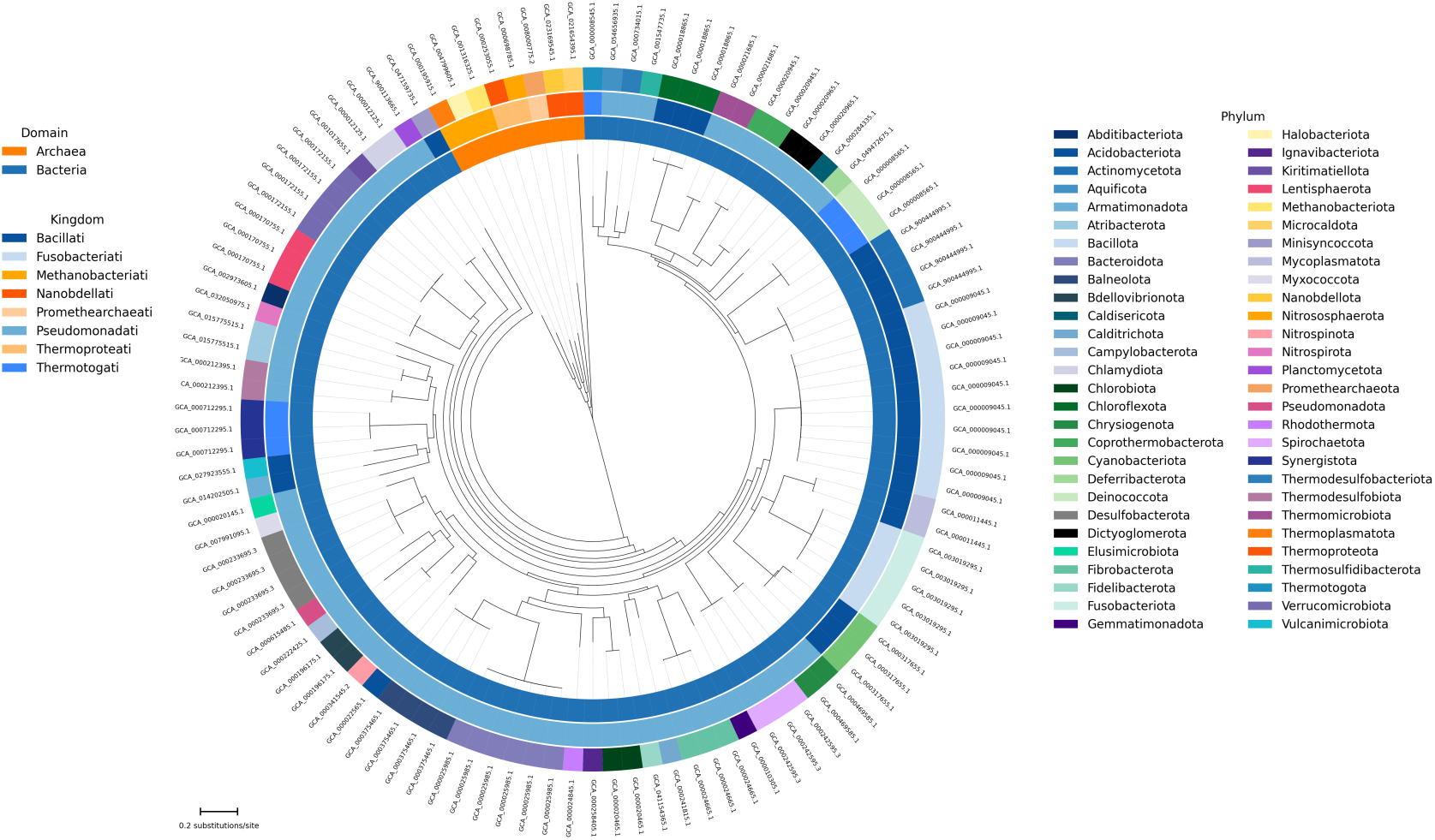
16S rRNA gene tree of 56 prokaryotic distinct phyla representatives. Full-length 16S rRNA gene sequences were identified from genome assemblies using RNAmmer, aligned with MAFFT, and used to infer a maximum-likelihood tree with IQ-TREE. The annotation rings show standardized NCBI taxonomy; from the outermost ring to the innermost ring, the colors indicate phylum, kingdom, and domain, respectively.

Repeated GCA labels in the circular tree correspond to multiple 16S rRNA copies retained from the same genome. **Supplementary Table 2** summarizes the ten genomes containing multiple retained copies within the reduced phylum-representative dataset used for **Figure 9**. This table is therefore a targeted description of the **Figure 9** tree dataset. It is not intended to represent the frequency or distribution of intragenomic heterogeneity across all 30,495 genomes shown in **Figure 4**. Among these ten representative genomes, the minimum within-genome 16S rRNA identity ranged from 95.26% to 99.87%. Two genomes had minimum identities below 99%: *Chloroflexus aurantiacus* at 98.98% and *Cyanobacterium stanieri* at 95.25%. It has been previously reported that for some genomes, different copies of the 16S rRNA genes can be significantly divergent [66]. For example, the ‘RiboGrove’ [69] website contains a list of the “top10” most divergent genomes.

**Supplementary Figure 4** shows a histogram of the “top ten” bacterial genomes from the RiboGrove list; note that for the type strain genomes (blue), all have >99% 16S identity (bar at the top of the histogram). The “Top 10 genomes” in green represent the genomes from the list of top 10 genomes with the largest 16S rRNA heterogeneity. Notice that for these ‘top10’ genomes in green, the internal lowest match for 16S rRNA from within the same genome ranges from less than 70% to about 85%. According to traditional 16S typing, these should belong to different phyla, classes, or orders (see **Figure 1**), not the same species, but these are all from the same genome. For 9 of the 10 genomes, there is a corresponding type strain for the species (shown in blue). For one genome (*Synechococcus* sp. NB0720_010) there is no type strain (the type strain for the *Synechococcus* genus (*S. elongatus*) is shown in light blue. The *Caminibacter mediatlanticus* type strain is also shown in light blue, as both genomes contain only a single rRNA gene, so the 16S rRNA % identity is to the corresponding sequence in the ribosomal database project (RDP [70]).

The analyses shown in **Figure 4**, **Supplementary Table 2**, and **Supplementary Figure 4** address different datasets and should be interpreted separately. Figure 4 reports the copy-number distribution across all 30,495 type-strain genomes and includes all qualifying full-length 16S rRNA copies. **Supplementary Table 2** is restricted to the ten multi-copy genomes occurring within the reduced phylum-representative dataset used for Figure 9. **Supplementary Figure 4** represents a separate targeted comparison between the highly heterogeneous genomes highlighted by RiboGrove and corresponding type-strain or reference genomes.

The 16S rRNA and FastAAI trees both recovered the broad separation between Archaea and Bacteria but differed substantially in their internal topologies (**Supplementary Figure 5**). For the Archaea-outgroup-rooted comparison, the unrooted Robinson–Foulds distance was 86 of 106 possible splits, corresponding to a normalized RF distance of 0.811. The quartet comparison identified 154,534 discordant quartets among 367,290 total quartets, giving a normalized quartet distance of 0.421. Branch-length-aware agreement was stronger than exact topological agreement: the Mantel correlations between patristic-distance matrices were approximately Pearson *r* = 0.91 and Spearman ρ = 0.81, with permutation *P* = 0.001. Leaf-order optimization reduced tanglegram crossings from 696 to 121 without changing either tree topology, although many crossings remained within bacterial lineages. These results indicate that the two methods preserve a similar broad distance structure while differing in many specific branching relationships.

## 4. Discussion

In this study, we systematically evaluated four widely used measures of prokaryotic similarity—16S rRNA gene identity, average nucleotide identity (ANI), FastAAI, and Mash distance—across a comprehensive set of 30,495 bacterial and archaeal type-strain genomes, spanning taxonomic levels from species to domain. Our goal was to empirically quantify the correspondence between these metrics and traditional taxonomic assignments and to provide practical, empirically derived thresholds that can guide microbial classification and comparative studies.

Consistent with decades of taxonomic practice, 16S rRNA similarity remains informative for broad taxonomic delineation. At higher ranks such as class and above, 16S identity retains some discriminative power, reflecting deeply conserved sequence motifs that are useful for broad phylogenetic grouping [15, 71], although as **Figure 1** shows, 16S identity provides less definitive resolution at the levels of phylum, class, and order. Further, we confirm that 16S often lacks resolution at the species level and sometimes at the genus level, consistent with previous observations that closely related taxa may share nearly identical 16S sequences despite substantial genomic divergence [72, 73]. This limitation constrains the utility of 16S alone for fine-scale taxonomic assignments in genomically diverse groups.

The present results distinguish two separate limitations of 16S rRNA analysis. First, 5,925 genomes (19%) lacked a recoverable full-length sequence, causing 4,551 same-species comparisons (28%) to fail before an identity value could be calculated. Second, even when a usable sequence was present, identity distributions overlapped among species, genus, and higher ranks. Nevertheless, 97% of valid same-species 16S comparisons passed the empirical threshold. Thus, missing marker-gene data were the principal practical limitation in this dataset, whereas overlapping sequence identities represented the principal biological limitation for species-level discrimination.

Whole-genome measures such as ANI have emerged as the *de facto* standard for fine-scale taxonomic resolution. Our analyses support prior benchmarks indicating that species boundaries correspond to an ANI range of ∼95–96% [16, 43], and that higher taxonomic ranks systematically diverge from this threshold in predictable ways. ANI captures overall genomic similarity across both coding and non-coding regions, making it robust to gene content variation that is invisible to single-locus markers. Tools such as FastANI further streamline ANI computation for large datasets, with reported high concordance to alignment-based ANI and dramatically reduced computational cost [44].

Mash and other MinHash-based sketching methods offer a distinct approach: by randomly sampling genomes as small representative sketches, Mash can estimate pairwise genomic distances orders of magnitude faster than alignment-based methods [49]. Our results show that Mash distance correlates strongly with ANI at shallow taxonomic depths, enabling rapid species- and genus-level clustering with minimal compute. This scalability is especially valuable in the context of large genome collections, where exhaustive ANI computations remain costly both in terms of CPU hours and I/O overhead. Nonetheless, because Mash distances are derived from sketch similarity rather than full sequence alignment, their sensitivity can vary with sketch parameters and genome completeness. Having said that, we have previously shown that Mash can reliably resolve subspecies phylogroups in *E. coli*, even from raw read files in the Sequence Read Archive (SRA) [74].

Our phylogenetic tree comparison results suggest that 16S rRNA and FastAAI should be viewed as complementary rather than competing approaches for higher-level taxonomic investigation. Both methods consistently recovered the broad Archaea/Bacteria separation, indicating shared deep-scale signal, but they differed substantially in many internal relationships and in recovery of monophyletic groups. This pattern is expected because 16S rRNA reflects the history of a single highly conserved marker, whereas FastAAI summarizes genome-wide protein-level relatedness and may capture broader aspects of evolutionary divergence. The combination of high patristic-distance correlations and high RF and quartet distances is biologically informative. It indicates that 16S rRNA and FastAAI preserve broadly similar geometries of evolutionary distances while disagreeing on many individual internal branches. This distinction explains how both approaches can recover major domain-level structure yet yield different placements for phyla or kingdoms.

One important outcome of the comparison is that empirical thresholds are useful only when interpreted within the taxonomic range for which each method retains resolution. The approximately 98.65% 16S rRNA identity guideline remains informative [16], but the substantial overlap among rank distributions shown in Figure 1 prevents it from serving as a universal species boundary. FastANI most clearly reflected the established 95–96% ANI species range [47, 48], whereas its values are compressed and provide little resolution at deeper ranks. Mash provides an efficient approximation of close genome relationships but increasingly returns the maximum distance as the sketch overlap disappears. FastAAI/Jaccard retained a broader gradient across higher ranks, although its thresholds should likewise be treated as empirical guides rather than universal nomenclatural boundaries.

These findings dovetail with results from curated taxonomies such as the Genome Taxonomy Database (GTDB), where species clusters are defined based on consistent genome phylogeny and ANI thresholds, and higher taxonomic levels are normalized across the tree [47, 48]. Our thresholds echo the empirically observed clusters in GTDB and extend them to include actionable values for Mash distances. The Sankey analyses also show that taxonomic sampling is highly uneven. *Methanobacteriati* accounted for 84% of the archaeal genomes, and *Bacillati* plus *Pseudomonadati* accounted for 99% of the bacterial genomes. Consequently, global summary statistics and empirical thresholds are influenced disproportionately by well-sampled cultivable lineages. At the same time, the 54 officially recognized phyla accounted for 99.8% of all genomes, suggesting that a simplified nomenclatural framework can retain nearly the entire dataset while reducing the influence of sparsely represented candidate and unclassified labels.

Our comparisons highlight complementary strengths among the methods. When researchers are working with large datasets or require rapid all-versus-all similarity estimates, Mash provides an efficient first pass that captures broad genomic similarity and correlates well with ANI [49, 75]. After initial clustering, selected subgroups can be refined using ANI or FastANI to resolve closely related taxa with higher fidelity. 16S rRNA remains a useful taxonomic anchor when only marker gene data are available, such as in many microbiome surveys. Still, its limited resolution at fine scales should caution against overinterpretation in isolation.

Furthermore, incomplete or draft genomes—common in environmental genomics— pose challenges for all methods; however, alignment-free approaches like Mash may be less sensitive to missing data than full sequence alignment or ANI, which require pairwise coverage. Despite this, genomes that are too fragmented may still yield unreliable similarity estimates across methods, underscoring the need for quality filtering and consistent preprocessing.

This large-scale empirical evaluation has several limitations. The standardized assignments were based primarily on NCBI taxonomy, which reflects a mixture of historical classifications, ongoing curation, and occasional inconsistencies among records. Some disagreement between sequence-based measurements and assigned labels may therefore reflect taxonomic annotation or nomenclatural history rather than failure of the method itself. Type-strain sampling was also highly uneven across kingdoms, phyla, and species, and many candidate or unclassified phyla were represented by only one or a few genomes. Draft assemblies introduced missing marker genes and fragmented genome content, while the empirical thresholds may vary among highly recombinant or unusually divergent lineages. The reported values should therefore be treated as dataset-wide benchmarks rather than immutable taxonomic rules.

### Conclusions

This study benchmarks 16S rRNA identity, FastANI, Mash, and FastAAI across a large, standardized set of prokaryotic type-strain genomes to evaluate how marker-gene and genome-wide methods perform across taxonomic ranks. The results show that 16S rRNA remains valuable for historical continuity, broad taxonomic placement, and comparisons with decades of microbial taxonomy. Still, its practical utility is limited by missing or fragmented full-length copies in draft genomes—5,925 genomes lacked a recoverable full-length sequence in this dataset—as well as intragenomic copy-number variation and overlapping identity ranges among taxonomic ranks. FastANI provides strong resolution for species-level and near-species-level comparisons, but it is less suitable for deeper taxonomic relationships. Mash offers a rapid alignment-free screening approach for the same or similar species within large genome collections, although its distances require careful interpretation, especially across broader evolutionary scales. FastAAI adds an important genome-wide amino-acid perspective and is particularly useful for comparisons beyond the species boundary, where nucleotide-based similarity becomes less informative.

Together, these results support the use of complementary methods rather than reliance on a single metric. Type-strain genomes provide a taxonomically meaningful benchmark because they anchor validly named species, but even this curated dataset contains draft assemblies, missing marker genes, and uneven representation across taxa. Separating initial method failures from threshold-based failures further clarifies whether disagreement reflects missing data, assembly quality, method limitations, or true biological divergence. Overall, benchmarking 16S rRNA, FastANI, Mash, and FastAAI across type-strain genomes provides practical guidance for selecting appropriate tools at different taxonomic scales and highlights the need for transparent, rank-aware, and genome-quality-aware approaches in modern prokaryotic taxonomy.

## 5. Author statements

### 5.1 Author contributions

DU conceptualized and designed the project and outlined the initial manuscript. DU drafted the Introduction, Results, and Discussion sections, and compiled the references. DU and AA reviewed and finalized the manuscript. All authors contributed to the writing and read and approved the final manuscript.

The following authors contributed to the figures:

1. - DU
2. - RM, AA, VAB and DU
3. - MZB and DU
4. - AA, and DU
5. - MZB and DU
6. - RM and DU
7. - VAB, and DU
8. - VAB, AA, and DU
9. - AA and DU

### 5.2 Conflicts of interest

The authors declare no conflict of interest.

### 5.3 Funding information

This work was funded in part by the McCasland Foundation Professor endowment and start-up funds for DU, from the College of Veterinary Medicine at Oklahoma State University, and an NIH grant (R01 GM148886).

## Acknowledgements

A project at UAMS inspired this manuscript, started in 2017 by George Garrity, Charles Parker, Mike Robeson, Visanu Wanchai, Kaleb Abram, and DU.

## Notes

https://github.com/ArghavanAlisoltani/30k-type-strain-taxonomy-benchmark

### Competing Interest Statement

The authors have declared no competing interest.

