## Supplementary Material for "Taxonomic Resolution of 16S rRNA, FastANI, Mash, and FastAAI across 30,495 Prokaryotic Type-Strain Genomes"

**[1] Department of Physiological Sciences, College of Veterinary Medicine,**

**[2] Department of Biochemistry and Molecular Biology,**

**Oklahoma State University, Stillwater Oklahoma, 74078, USA**

**Supplementary Table 1.** Prokaryotic genera named after pioneer microbiologists.

| Year | Honoree | Genus | # Type Genomes | Notable Contribution |
|---|---|---|---|---|
| **1676** | Antonie van Leeuwenhoek | *Leeuwenhoekiella* | **10** | First described bacteria |
| **1867** | Joseph Lister | *Listeria* | **58** | Published the principles of antiseptic surgery. |
| **1872** | Ferdinand Cohn | *Cohnella* | **43** | Published an early bacterial classification system |
| **1879** | Albert Neisser | *Neisseria* | **66** | Discovered the gonorrhea bacterium. |
| **1880** | Louis Pasteur | *Pasteurella* | **13** | Proposed germ theory of disease. |
| **1884** | Hans Christian Gram | *Christiangramia* | **22** | Invented the Gram stain technique. |
| **1885** | Theodor Escherich | *Escherichia* | **26** | Discovered the bacterium we now know as *E. coli*. |
| **1885** | Daniel E. Salmon | *Salmonella* | **41** | Co-discovered *Salmonella*. |
| **1887** | David Bruce | *Brucella* | **67** | Identified Malta fever bacterium. |
| **1894** | Alexandre Yersin | *Yersinia* | **43** | Co-discovered the plague bacillus. |
| **1897** | Kiyoshi Shiga | *Shigella* | **11** | Discovered the dysentery bacillus. |
| **1898** | Martinus Beijerinck | *Beijerinckia* | **2** | Described the first virus and pioneered environmental microbiology. |
| **1906** | Jules Bordet | *Bordetella* | **30** | Co-discovered the whooping cough bacillus (*Bordetella pertussis*). |
| **1969** | Thomas Brock | *Brockia* | **1** | Discovered high-temperature extremophiles like *Thermus aquaticus*. |
| **1977** | Carl Woese | *Woeseia* | **1** | Discovered Archaea as a distinct third domain of life. |
| **1981** | Karl Stetter | *Stetteria* | **1** | Pioneered the discovery of hyperthermophilic Archaea. |

**Supplementary Table 2**. Pairwise 16S rRNA identity statistics for 10 selected genomes with multiple 16S rRNA copies.

| **ID** | **species** | **phylum** | **16S copy count** | **Min within genome identity %** | **Max within genome identity %** |
|---|---|---|---|---|---|
| GCA_000008565.1 | Deinococcus radiodurans | Deinococcota | 3 | 99.8658 | 100 |
| GCA_000009045.1 | Bacillus subtilis | Bacillota | 10 | 99.3485 | 100 |
| GCA_000011445.1 | Mycoplasma mycoides | Mycoplasmatota | 2 | 99.4702 | 99.4702 |
| GCA_000018865.1 | Chloroflexus aurantiacus | Chloroflexota | 3 | 98.9817 | 99.8643 |
| GCA_000020945.1 | Coprothermobacter proteolyticus | Coprothermobacterota | 2 | 99.4751 | 99.4751 |
| GCA_000024665.1 | Fibrobacter succinogenes | Fibrobacterota | 3 | 99.3275 | 99.5965 |
| GCA_000170755.1 | Lentisphaera araneosa | Lentisphaerota | 3 | 99.867 | 99.9335 |
| GCA_000172155.1 | Verrucomicrobium spinosum | Verrucomicrobiota | 4 | 99.869 | 100 |
| GCA_000317655.1 | Cyanobacterium stanieri | Cyanobacteriota | 3 | 95.2575 | 99.3902 |
| GCA_000469585.1 | Chrysiogenes arsenatis | Chrysiogenota | 2 | 99.8753 | 99.8753 |

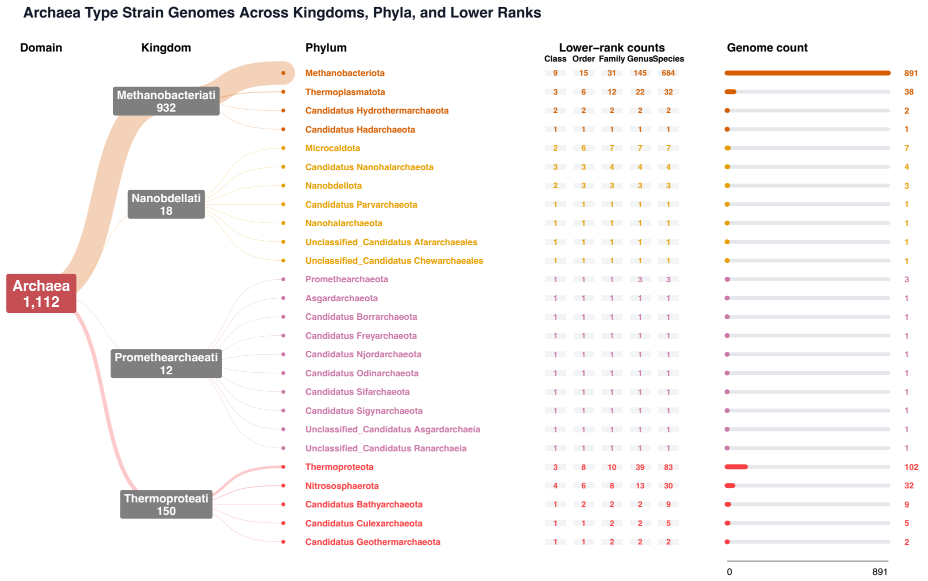


**Supplementary Figure 1A. Taxonomic distribution of 1,112 archaeal type-strain genomes.** Sankey flows connect the archaeal domain to four displayed kingdom groups and 26 phyla. For each phylum, the adjacent columns report the numbers of unique classes, orders, families, genera, and species, together with the total number of genomes. Flow widths and horizontal bars are proportional to genome counts.

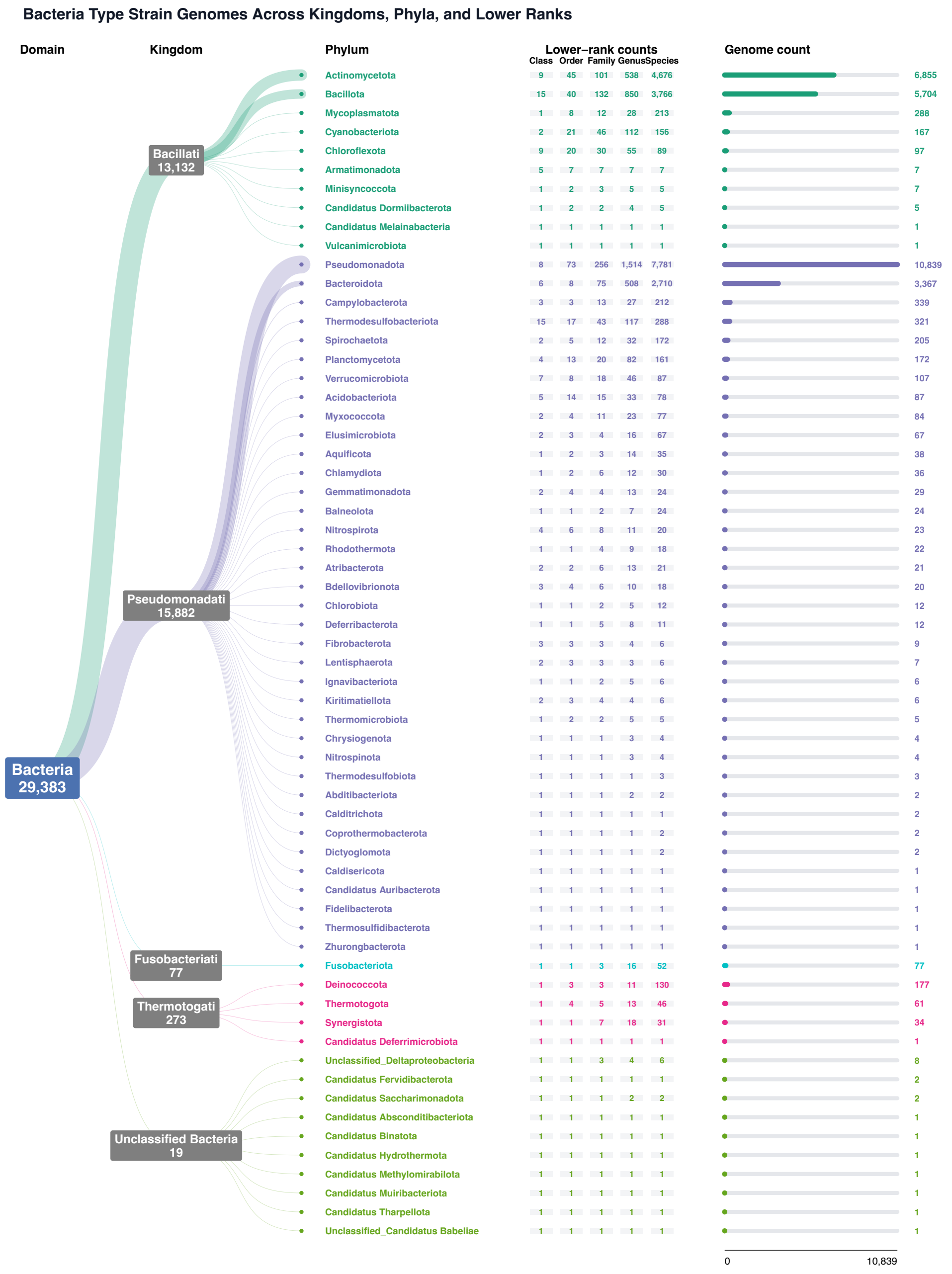


**Supplementary Figure 1B. Taxonomic distribution of 29,383 bacterial type-strain genomes.** Sankey flows connect the bacterial domain to five displayed kingdom-level groups and 62 phyla. For each phylum, the adjacent columns report the numbers of unique classes, orders, families, genera, and species, together with the total number of genomes. Unclassified Bacteria combines 19 genomes assigned to two unclassified bacterial taxonomy labels. Flow widths and horizontal bars are proportional to genome counts.

### A.Carl Woese Original 12 Groups –> Modern Phyla

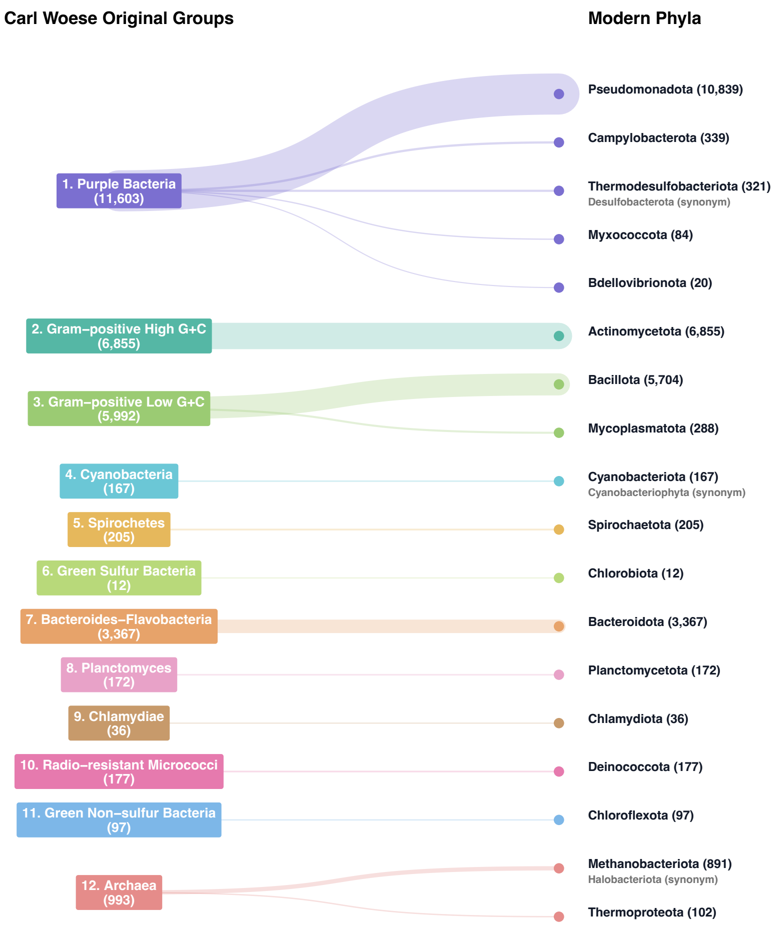

B. New Phyla Outside the Original Woese Groups

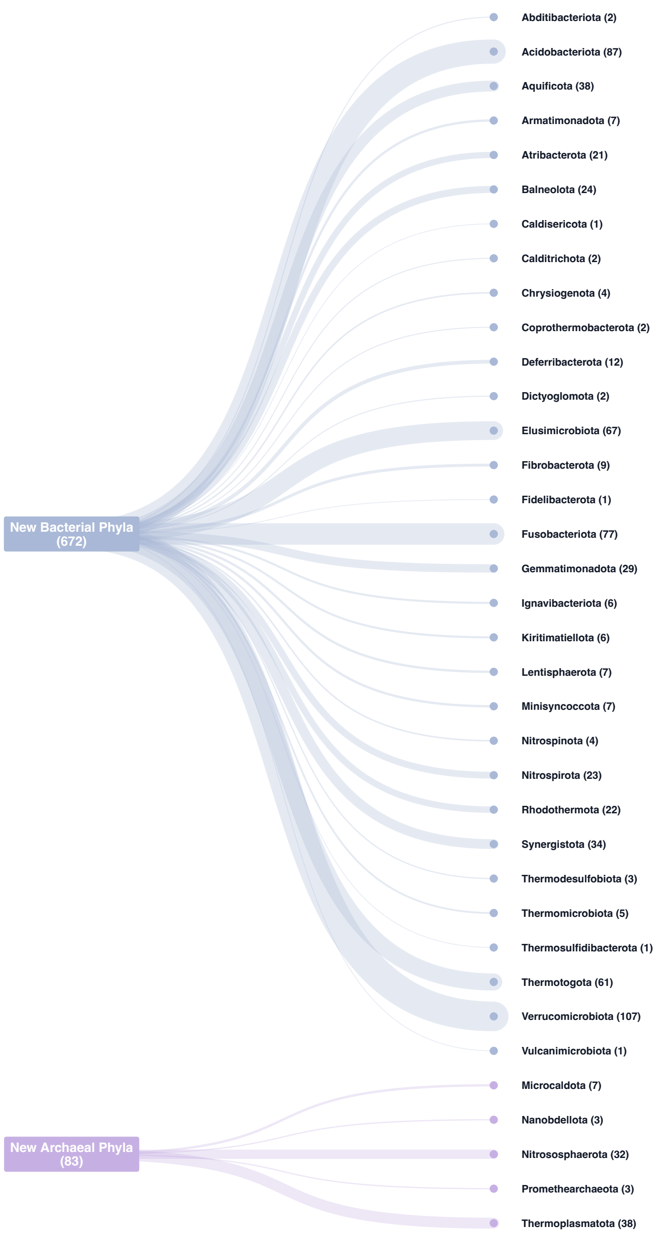


**Supplementary Figure 2. Relationship between the original Woese groups and currently recognized prokaryotic phyla.** Panel A maps the 12 original Woese groups to 18 corresponding modern phyla containing 29,676 genomes. Panel B shows 36 additional officially recognized phyla containing 755 genomes. Together, the 54 recognized phyla contain 30,431 of the 30,495 type-strain genomes (99.8%). Numbers in parentheses indicate genome counts; the remaining 64 genomes were assigned to candidate, unclassified, or other phylum labels not included in these panels.

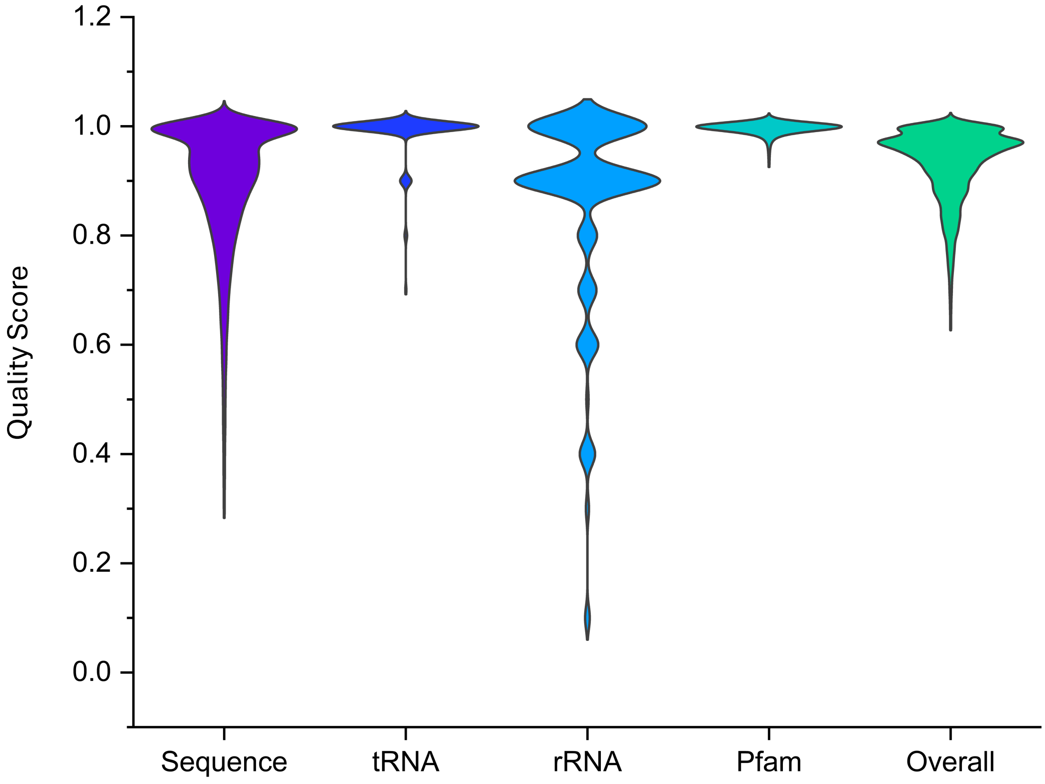


**Supplementary Figure 3. Genome-quality score distributions across all 30,495 type-strain genomes.** Violin plots show the sequence, tRNA, rRNA, Pfam essential-gene, and overall quality-score distributions. Higher values indicate greater estimated completeness or quality.

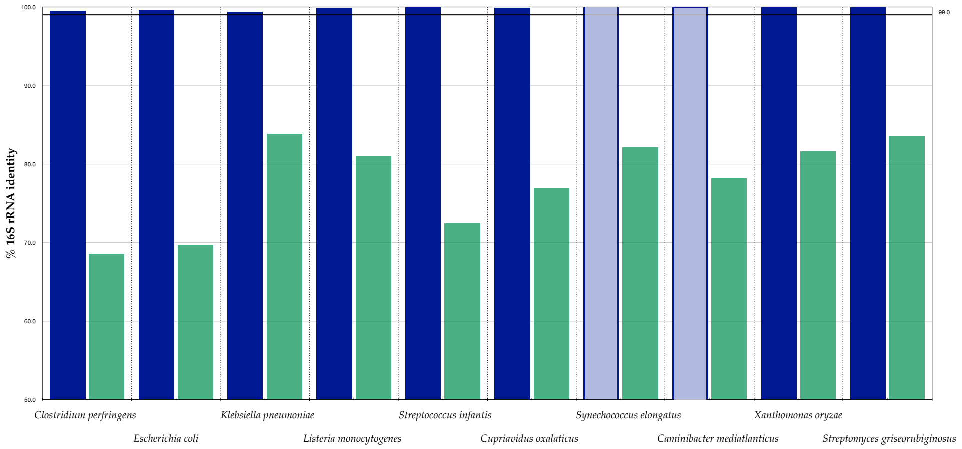


**Supplementary Figure 4. Comparison of intragenomic 16S rRNA heterogeneity in selected public genomes and corresponding type-strain references.** Green bars show the minimum within-genome 16S rRNA identity for the ten highly heterogeneous genomes highlighted by RiboGrove. Dark-blue bars show the corresponding values for available type-strain genomes. Light-blue bars indicate references for which only one 16S rRNA copy was recovered; in these cases, identity was calculated against the corresponding Ribosomal Database Project sequence. The horizontal line marks 99% identity.

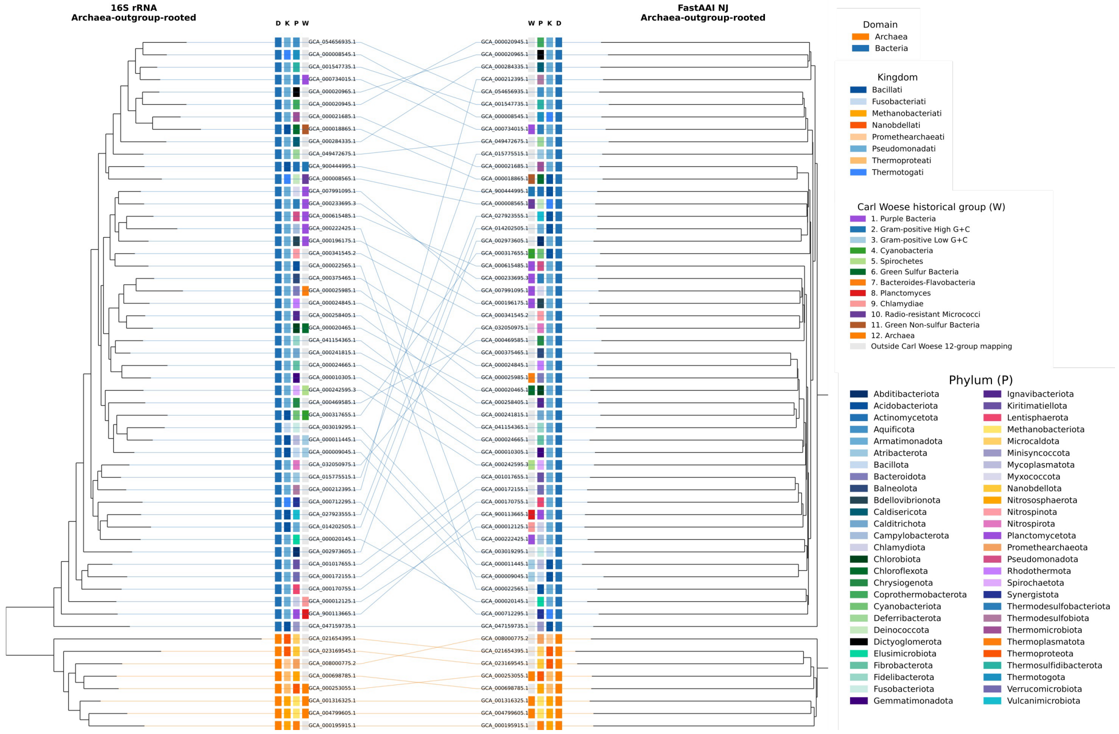


**Supplementary Figure 5** Tanglegram comparing 16S rRNA and FastAAI phylogenies for 56 prokaryotic genomes, both rooted with Archaea as the outgroup. The left tree is the 16S rRNA tree and the right is the FastAAI neighbour-joining tree. Connecting lines join the same genome in the two trees and are colored by domain (orange = Archaea, blue = Bacteria); tip labels (GCA accessions) are shown in black. Four metadata strips flank each tree — D = domain, K = kingdom, P = phylum (official phylum), W = Carl Woese's historical group — colored according to the legends at right.
